# Whole organism lineage tracing by combinatorial and cumulative genome editing

**DOI:** 10.1101/052712

**Authors:** Aaron McKenna, Gregory M. Findlay, James A. Gagnon, Marshall S. Horwitz, Alexander F. Schier, Jay Shendure

## Abstract

Multicellular systems develop from single cells through a lineage, but current lineage tracing approaches scale poorly to whole organisms. Here we use genome editing to progressively introduce and accumulate diverse mutations in a DNA barcode over multiple rounds of cell division. The barcode, an array of CRISPR/Cas9 target sites, records lineage relationships in the patterns of mutations shared between cells. In cell culture and zebrafish, we show that rates and patterns of editing are tunable, and that thousands of lineage-informative barcode alleles can be generated. We find that most cells in adult zebrafish organs derive from relatively few embryonic progenitors. Genome editing of synthetic target arrays for lineage tracing (GESTALT) will help generate large-scale maps of cell lineage in multicellular systems.

## Introduction

The tracing of cell lineages was pioneered in nematodes by Charles Whitman in the 1870s, at a time of controversy surrounding Ernst Haeckel’s theory of recapitulation (1). This line of work culminated a century later in the complete description of mitotic divisions in the roundworm *C. elegans*- a tour de force facilitated by its visual transparency as well as the modest size and invariant nature of its cell lineage (2).

Over the past century, a variety of creative methods have been developed for tracing cell lineage in developmentally complex organisms (3). In general, subsets of cells are marked and their descendants followed as development progresses. The ways in which cell marking has been achieved include dyes and enzymes (4-6), cross-species transplantation (7), recombinase-mediated activation of reporter gene expression (8, 9), insertion of foreign DNA (10-12), and naturally occurring somatic mutations (13-15). However, despite many powerful applications, these methods have limitations for the large-scale reconstruction of cell lineages in multicellular systems. For example, dye and reporter gene-based cell marking are uninformative with respect to the lineage relationships *between* descendent cells. Furthermore, when two or more cells are independently but equivalently marked, the resulting multitude of clades cannot be readily distinguished from one another. Although these limitations can be overcome in part with combinatorial labeling systems (16, 17) or through the introduction of diverse DNA barcodes (10-12), these strategies fall short of a system for inferring lineage relationships throughout an organism and across developmental time. In contrast, methods based on somatic mutations have this potential, as they can identify lineages and sub-lineages within single organisms (13, 18). However, somatic mutations are distributed throughout the genome, necessitating whole genome sequencing, (14, 15), which is expensive to scale beyond small numbers of cells and not readily compatible with *in situ* readouts (19, 20).

What are the requirements for a system for comprehensively tracing cell lineages in a complex multicellular system? First, it must uniquely and incrementally mark cells and their descendants over many divisions and in a way that does not interfere with normal development. Second, these unique marks must accumulate irreversibly over time, allowing the reconstruction of lineage trees. Finally, the full set of marks must be easily read out in each of many single cells.

We hypothesized that genome editing, which introduces diverse, irreversible edits in a highly programmable fashion (21), could be repurposed for cell lineage tracing in a way that realizes these requirements. To this end, we developed <u>g</u>enome <u>e</u>diting of <u>s</u>ynthetic <u>t</u>arget <u>a</u>rrays for <u>l</u>ineage <u>t</u>racing (GESTALT), a method that uses CRISPR/Cas9 genome editing to accumulate combinatorial sequence diversity to a compact, multi-target, densely informative barcode. Importantly, edited barcodes can be efficiently queried by a single sequencing read from each of many single cells (Fig. 1A). In both cell culture and in the zebrafish *Danio rerio*, we demonstrate the generation of thousands of uniquely edited barcodes that can be related to one another to reconstruct cell lineage relationships. In adult zebrafish, we observe that the majority of cells of each organ are derived from a small number of progenitor cells. Furthermore, ancestral progenitors, inferred on the basis of shared edits amongst subsets of derived alleles, make highly non-uniform contributions to germ layers and organ systems.

**Fig. 1.**
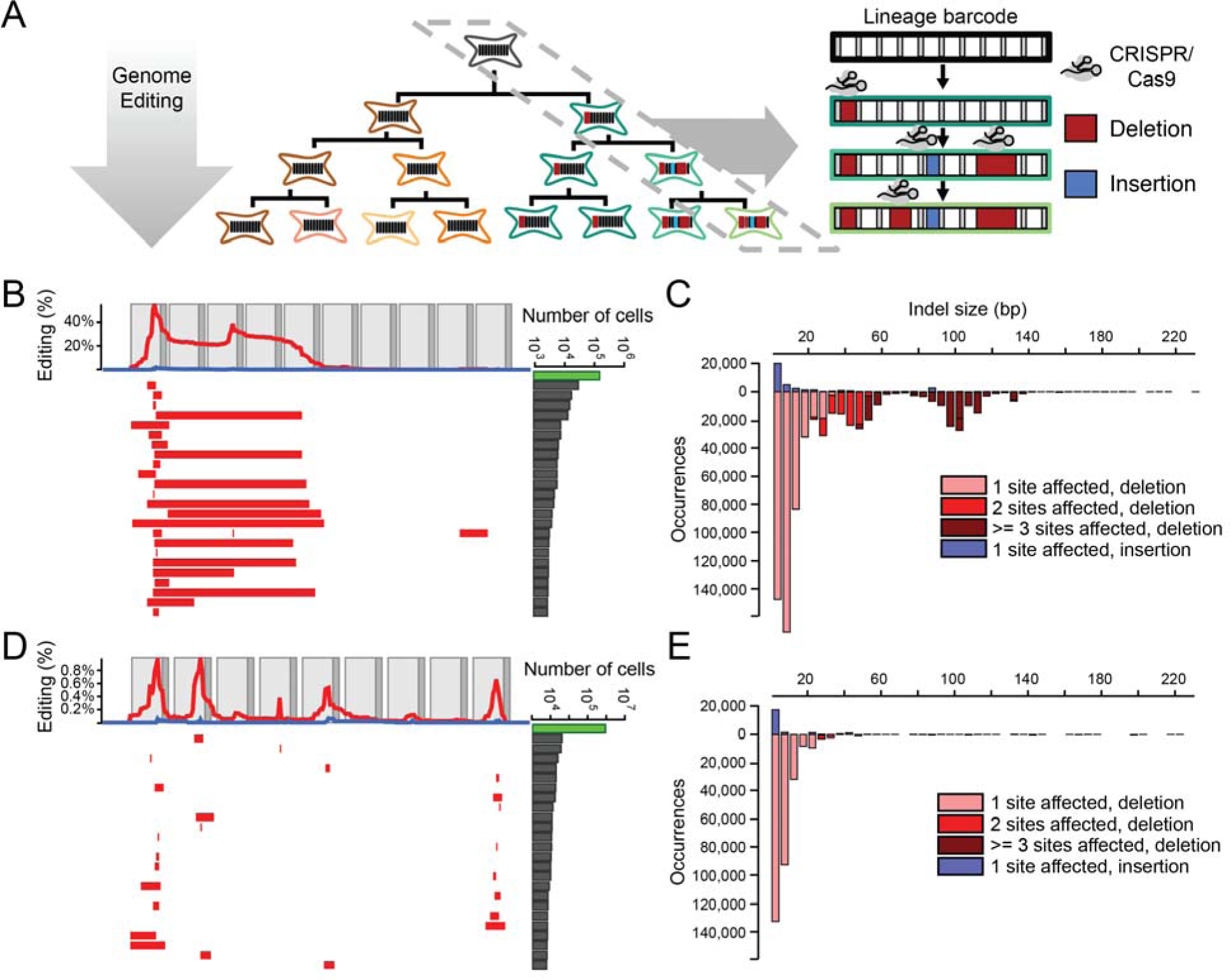
Genome editing of synthetic target arrays for lineage tracing (GESTALT). (**A**) An unmodified array of CRISPR/Cas9 target sites (*i.e.*, a barcode) is engineered into a genome. Editing reagents are introduced during expansion of cell culture or *in vivo* development of an organism, such that distinct lineages stably accumulate different edits to the barcode. Because the resulting edited barcodes are compact, derivative sequences present within a population of cells can be queried by PCR and sequencing, such that each sequencing read corresponds to the barcode derived from a single cell. Edited barcodes that are allelic, *i.e.* identical in sequence, but differ by UMI are inferred to have derived from distinct but closely related cells. The lineage relationships of alleles that differ in sequence can often be inferred on the basis of editing patterns. (**B**) The 25 most frequent alleles from the edited v1 barcode are shown. Each row corresponds to a unique sequence, with red bars indicating deleted regions and blue bars indicating insertion positions. Blue bars begin at the insertion site, with their width proportional to the size of the insertion, which will rarely obscure immediately adjacent deletions. The number of reads observed for each allele is plotted at the right (log10 scale; the green bar corresponds to the unedited allele). The frequency at which each base is deleted (red) or flanks an insertion (blue) is plotted at the top. Light gray boxes indicate the location of CRISPR protospacers while dark gray boxes indicate PAM sites. For the v1 array, inter-target deletions involving sites 1, 3 and 5, or focal (single target) edits of sites 1 and 3 were observed predominantly. (**C**) A histogram of the size distribution of insertion (top) and deletion (bottom) edits to the v1 array is shown. The colors indicate the number of target sites impacted. Although most edits are short and impact a single target, a substantial proportion of edits are inter-target deletions. (**D**) We tested three array designs in addition to v1, each comprising nine to ten weaker off-target sites for the same sgRNA (v2-v4) (22). Editing of the v2 array is shown with layout as described in panel (B). Editing of the v3 and v4 array are shown in fig. S3, A and B. The weaker sites within these alternative designs exhibit lower rates of editing than the v1 array, but also a much lower proportion of inter-target deletions. (**E**) A histogram of the size distribution of insertion (top) and deletion (bottom) edits to the v2 array is shown. In contrast with the v1 array, almost all edits impact only a single target.

## Results

### Combinatorial and cumulative editing of a compact genomic barcode in cultured cells

To test whether genome editing can be used to generate a combinatorial diversity of mutations within a compact region, we synthesized a contiguous array of ten CRISPR/Cas9 targets (protospacers plus PAM sequences) separated by 3 base-pair (bp) linkers (total length of 257 bp). The first target perfectly matched one single guide RNA (sgRNA), while the remainder were off-target sites for the same sgRNA, ordered from highest to lowest activity (22). This array of targets (‘v1 barcode’) was cloned downstream of an EGFP reporter in a lentiviral construct (23). We then transduced HEK293T cells with lentivirus and used FACS to purify an EGFP-v1 positive population. To edit the barcode, we co-transfected these cells with a plasmid expressing Cas9 and the sgRNA and a vector expressing DsRed. Cells were sorted three days post-transfection for high DsRed expression, and genomic DNA (gDNA) was harvested on day 7. The v1 barcode was PCR amplified and the resulting amplicons subjected to deep sequencing.

To minimize confounding sequencing errors, which are primarily substitutions, we analyzed edited barcodes for only insertion-deletion changes relative to the ‘wild-type’ v1 barcode. In this first experiment, we observed 1,650 uniquely edited barcodes (each observed in ≥25 reads) with diverse edits concentrated at the expected Cas9 cleavage sites, predominantly inter-target deletions involving sites 1, 3 and 5, or focal edits of sites 1 and 3 (Fig. 1, B and C, and table S1). These results show that combinatorial editing of the barcode can give rise to a large number of unique sequences, *i.e.* “alleles”.

To evaluate reproducibility, we transfected the same editing reagents to cultures expanded from three independent EGFP-v1 positive clones. Targeted RT-PCR and sequencing of EGFP-v1 RNA showed similar distributions of edits to the v1 barcode in the transcript pool, between replicates as well as in comparison to the previous experiment (fig. S1). These results show that the observed editing patterns are largely independent of the site of integration and that edited barcodes can be queried from either RNA or DNA.

To evaluate how editing outcomes vary as a function of Cas9 expression, we co-transfected EGFP-v1 positive cells with a plasmid expressing Cas9 and the sgRNA as well as an DsRed vector, and after four days sorted cells into low, medium, and high DsRed bins and harvested gDNA. Overall editing rates matched DsRed expression (frequency of non-wild-type barcodes: low DsRed = 40%; medium DsRed = 69%; high DsRed = 91%). The profile of edits observed remained similar, but there were fewer inter-target deletions in the lower DsRed bins (fig. S2). These results show that adjusting expression levels of editing reagents can be used to modify the rates and patterns of barcode editing.

We also synthesized and tested three barcodes (v2-v4) with nine or ten weaker off-target sites for the same sgRNA as used for v1 (22). Genome editing resulted in derivative barcodes with substantially fewer edits than seen with the v1 barcode, but a much greater proportion of these edits were to a single target site, *i.e.* fewer inter-target deletions were observed (Fig. 1, D and E, and fig. S3, A and B). As only a few targets were substantially edited in designs v1-v4, we combined the most highly active targets to a new, twelve target barcode (v5). This barcode exhibited more uniform usage of constituent targets, but with relative activities still ranging over two orders of magnitude (fig. S3C and table S1). These results illustrate the potential value of iterative barcode design.

To determine whether the means of editing reagent delivery influences patterns of barcode editing, we introduced a lentiviral vector expressing Cas9 and the same sgRNA to cells containing the v5 barcode (24). After two weeks of culturing a population bottlenecked to 200 cells by FACS, we observed diverse barcode alleles but with substantially fewer inter-target deletions than with episomal delivery of editing reagents (fig. S3D). This finding demonstrates that the allelic spectrum can also be modulated by the delivery mode of editing reagents.

Taken together, these results show that editing multiple target sites within a compact barcode can generate a combinatorial diversity of alleles, and also that these alleles can be read out by single sequencing reads derived from either DNA or RNA. Rates and patterns of barcode editing are tunable by using targets with different activities and/or off-target sequences, by iteratively recombining targets to new barcode designs, and by modulating the concentration and means of delivery of editing reagents.

### Reconstruction of lineage relationships in cultured cells

To determine whether GESTALT could be used to reconstruct lineage relationships, we applied it to a designed lineage in cell culture (Fig. 2). A monoclonal population of EGFP-v1 positive cells was transfected with editing reagents to induce a first round of mutations in the v1 barcode. Clones derived from single cells were expanded, sampled, split, and re-transfected with editing reagents to induce a second round of mutations of the v1 barcode. For each clonal population, two 100-cell samples of the re-edited populations were expanded and harvested for gDNA. In these experiments, we began incorporating unique molecular identifiers (UMIs; 10 bp) during amplification of barcodes by a single round of polymerase extension (fig. S4A). Each UMI tags the single barcode present within each single cell, thereby allowing for correction of subsequent PCR amplification bias and enabling each UMI-barcode combination to be interpreted as deriving from a single cell (25).

**Fig. 2.**
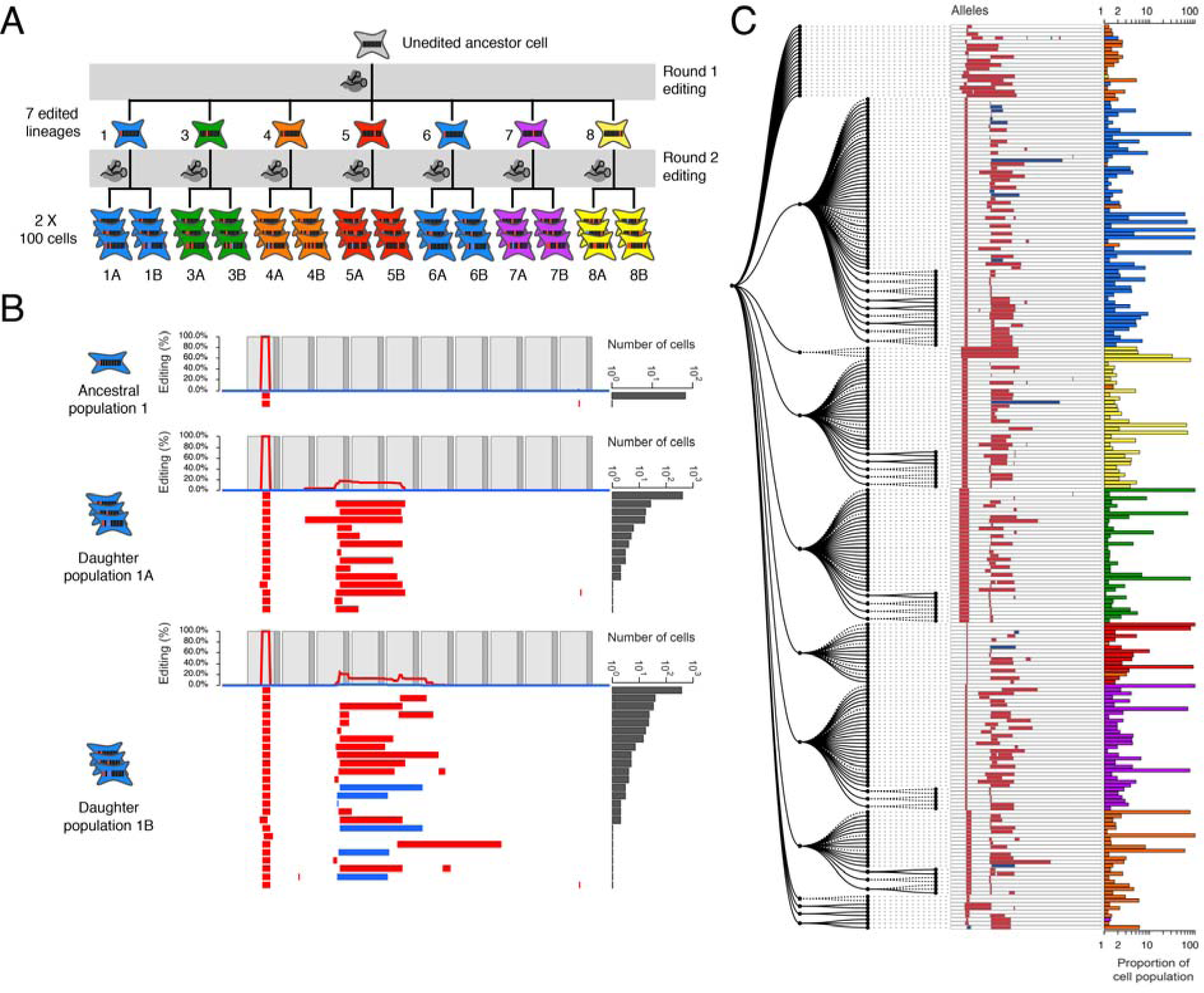
Reconstruction of a synthetic lineage based on genome editing and targeted sequencing of edited barcodes. (**A**) A monoclonal population of cells was subjected to editing of the v1 array. Single cells were expanded, sampled (#1 to #12), re-transfected to induce a second round of barcode editing, and then expanded and sampled from 100-cell subpopulations (#1a, 1b to #12a, 12b). For clarity, the five clones where the original population was unedited are not shown. (**B**) Alleles observed in the synthetic lineage experiment are shown, with layout as described in the Fig. 1B legend. #1 represents sampling of cells that had been subjected to only the first round of editing; virtually all cells contain a shared edit to the first target. #1a and #1b are derived from #1 but subjected to a second round of editing prior to sampling. These retain the edit to the first target, but subpopulations bear additional edits to other targets. (**C**) Parsimony (fig. S4B) using all alleles from the seven cell lineages represented in panel (A). Clade membership and abundance of each allele are shown on the right. #4 appears to be derived from two cells, one edited and the other wild-type: only 62% of lineage #4 falls into a single clade, consistent with the proportion (64%) of the lineage determined by sequencing to be edited after the first round. It is assumed that cells unedited in the first round either accrued edits matching other lineages (thus causing mixing), or accrued different edits (thus remaining outside the major clades).

Seven of twelve clonal populations we isolated contained mutations in the v1 barcode that were unambiguously introduced during the first round of editing (Fig. 2A). Additional edits accumulated in re-edited cells but generally did not disrupt the early edits (Fig. 2B and fig. S5). We next sought to reconstruct the lineage relationships between all alleles observed in the experiment using a maximum parsimony approach (fig. S4B)(26). The resultant tree contained major clades that were defined by the early edits present in each lineage (Fig. 2C). Four clonal populations (#3, #5, #7 and #8) were cleanly separated upon lineage reconstruction, with >99.7% of cells accurately placed into each lineage’s major clade. Two lineages (#1 and #6) were mixed because they shared identical mutations from the first round of editing. These most likely represent the recurrence of the same editing event across multiple lineages, but could also have been daughter cells subsequent to a single, early editing event prior to isolating clones. Consequently, 99.9% of cells of these two lineages were assigned to a single clade (Fig. 2C, blue). One clonal population (#4) appears to have derived from two independent cells, one of which harbored an unedited barcode. Later editing of these barcodes confounded the assignment of this lineage on the tree. Overall, however, these results demonstrate that GESTALT can be used to capture and reconstruct cell lineage relationships in cultured cells.

### Combinatorial and cumulative editing of a compact genomic barcode in zebrafish

To test the potential of GESTALT for *in vivo* lineage tracing in a complex multicellular organism, we turned to the zebrafish *Danio rerio*. We designed two new barcodes, v6 and v7, each with ten sgRNA target sites that are absent from the zebrafish genome and predicted to be highly editable (methods). In contrast to v1-v5, in which the target sites are variably editable by one sgRNA, the targets within v6 or v7 are designed to be edited by distinct sgRNAs. We generated transgenic zebrafish that harbor each barcode in the 3’ UTR of DsRed driven by the ubiquitin promoter (27, 28) and a GFP marker that is expressed in the cardiomyocytes of the heart (fig. S6) (29). To evaluate whether diverse alleles could be generated by *in vivo* genome editing, we injected Cas9 and ten different sgRNAs with perfect complementarity to the barcode target sites into single-cell v6 embryos (Fig. 3A). Editing of integrated barcodes had no noticeable effects on development (fig. S7). To characterize barcode editing *in vivo*, we extracted gDNA from a series of single 30 hours post fertilization (hpf) embryos, and UMI-tagged, amplified and sequenced the v6 barcode. In control embryos (Cas9-; n = 2), all 4,488 captured barcodes were unedited. In contrast, in edited embryos (Cas9+; n = 8), fewer than 1% of captured barcodes were unedited. We recovered barcodes from hundreds of cells per embryo (median 943; range 257−2,832) and identified dozens to hundreds of alleles per embryo (median 225; range 86-1,323). 41% +/− 10% of alleles were observed recurrently within single embryos, most likely reflecting alleles that were generated in a progenitor of two or more cells. Fewer than 0.01% of alleles were shared in pairwise comparisons of embryos, revealing the highly stochastic nature of editing in different embryos. These results demonstrate that GESTALT can generate very high allelic diversity *in vivo*.

**Fig. 3.**
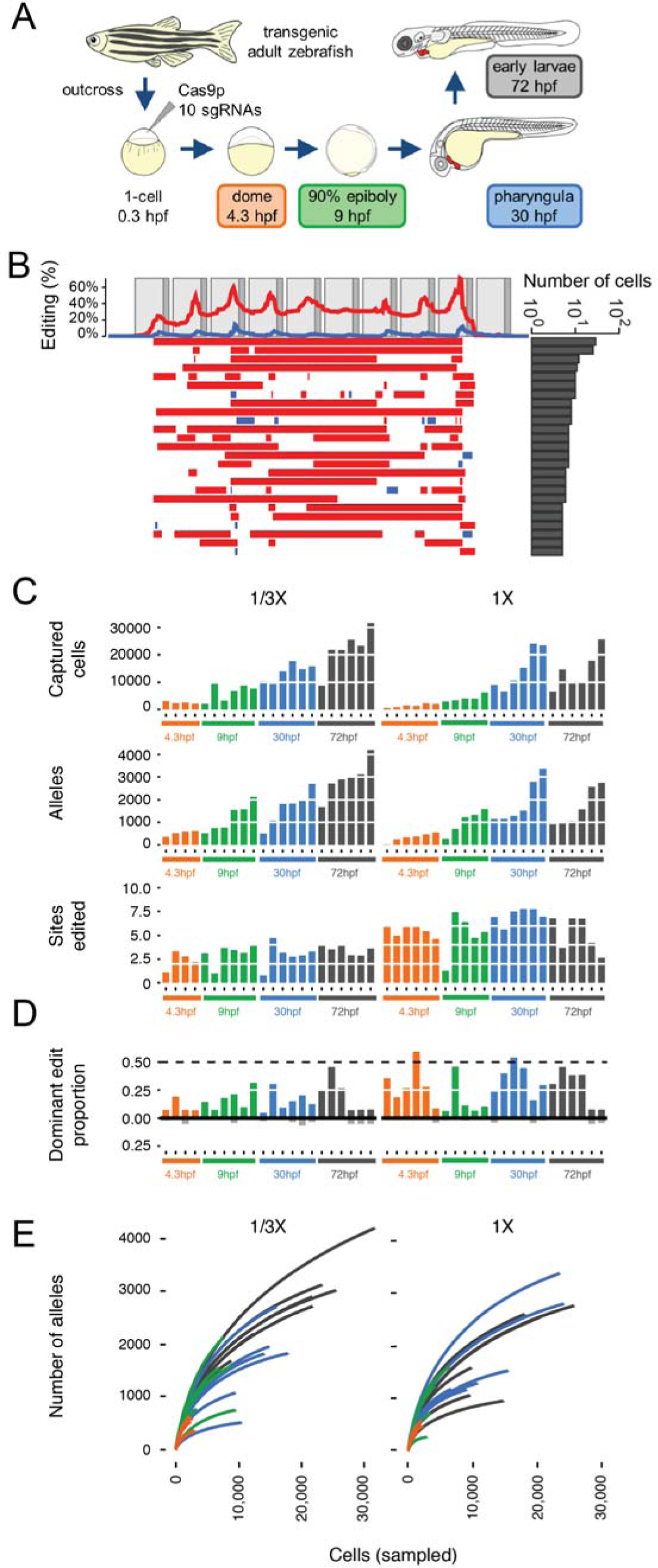
Generating combinatorial barcode diversity in transgenic zebrafish. (**A**) One-cell zebrafish embryos were injected with complexed Cas9 ribonucleoproteins (RNPs) containing sgRNAs that matched each of the 10 targets in the array (v6 or v7). Embryos were collected at time points indicated. UMI-tagged barcodes were amplified and sequenced from genomic DNA. (**B**) Patterns of editing in alleles recovered from a 30 hpf v6 embryo, with layout as described in the Fig. 1B legend. (**C**) Bar plots show the number of cells sampled (top), unique alleles observed (middle) and proportion of sites edited (bottom) for 45 v7 embryos collected at four developmental time-points and two levels of Cas9 RNP (1/3x, 1x). Colors correspond to stages shown in panel (A). Although more alleles are observed with sampling of larger numbers of cells at later time points, the proportion of target sites edited remains relatively constant. (**D**) Bar plots show the proportion of edited barcodes containing the most common editing event in a given embryo. Six of 45 embryos had the most common edit in approximately 50% of cells (dashed line), consistent with this edit having occurred at the two-cell stage (see fig. S8A for example). Colors correspond to stages shown in panel (A). These same edits are rarer or absent in other embryos (black bars below). (**E**) For each of the 45 v7 embryos, all barcodes observed were sampled without replacement. The cumulative number of unique alleles observed as a function of the number of cells sampled is shown (average of the 100 iterations shown per embryo; two levels of Cas9 RNP: 1/3x on left, 1x on right). The number of unique alleles observed, even in later stages where we are sampling much larger numbers of cells, appears to saturate, and there is no consistent pattern supporting substantially greater diversity in later time-points, consistent with the bottom row of panel (C) in supporting the conclusion that the majority of editing occurs before dome stage.

### Reconstruction of lineage relationships in embryos

To evaluate whether lineage relationships can be reconstructed using edited barcodes, we focused on the v6 embryo with the lowest rates of inter-target deletions and edited target sites (Fig. 3B; avg. 58% +/− 27% of target sites no longer a perfect match to the unedited target, compared to 87% +/− 21% for all other 30 hpf v6 embryos). Application of our parsimony approach (fig. S4B) to the 1,961 cells in which we observed 1,323 distinct alleles generated the large tree shown in Fig. 4. 1,307 of the 1,323 (98%) alleles could be related to at least one other allele by one or more shared edits, 85% by two or more shared edits, and 56% by three or more shared edits. These results illustrate the principle of using patterns of shared edits between distinct barcode alleles to reconstruct their lineage relationships *in vivo*.

**Figure 4.**
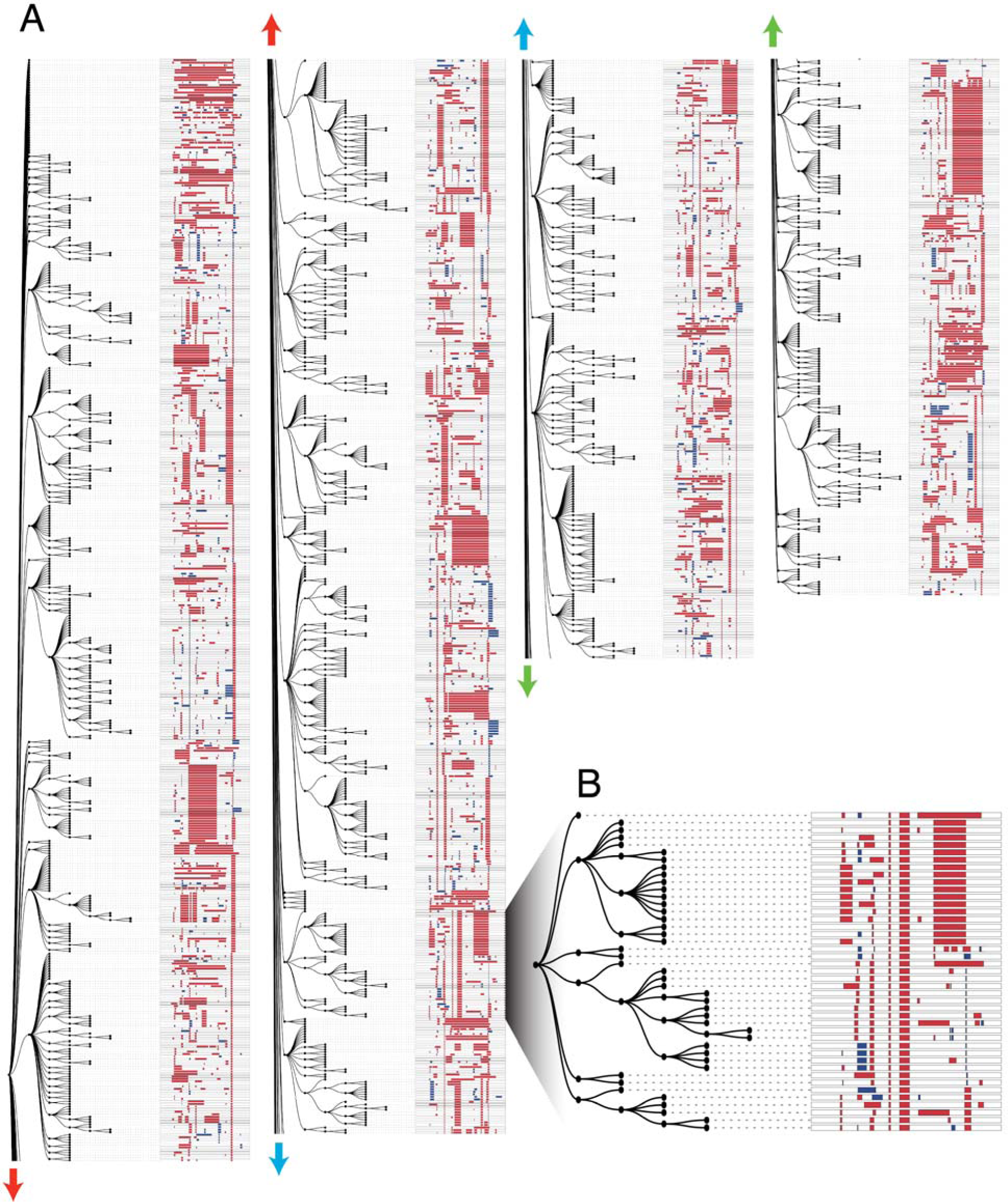
Lineage reconstruction of an edited zebrafish embryo. (**A**) A lineage reconstruction of 1,323 alleles recovered from the v6 embryo also represented in Fig. 3B, generated by a maximum parsimony approach (fig. S4B). A dendrogram to the left of each column represents the lineage relationships, and the alleles are represented at right. Each row represents a unique allele. Matched colored arrows connect subsections of the tree together. There are many large clades of alleles sharing specific edits, as well as sub-clades defined by ‘dependent’ edits. These dependent edits occur within a clade defined by a more frequent edit but are rare or absent elsewhere in the tree. (**B**) A portion of the tree is shown at higher resolution. A single edit is shared by all alleles in this clade. Six independent edits define descendent sub-clades within this clade, and further edits define additional sub-sub-clades within the clade.

### Developmental timing of barcode editing

To determine the developmental timing of barcode editing, we injected Cas9 and ten sgRNAs into one-cell stage v7 transgenic embryos and harvested genomic DNA before gastrulation (dome stage, 4.3 hpf; n = 10 animals), after gastrulation (90% epiboly / bud stage, 9 hpf; n = 11 animals), at pharyngula stage (30 hpf; n = 12 animals), and from early larvae (72 hpf; n = 12 animals) (Fig. 3A). We recovered barcode sequences from a median of 8,785 cells per embryo (range 461-31,640; total of 45 embryos), comprising a median of 1,223 alleles per embryo (range 15-4,195) (Fig. 3C). Within single embryos, 65% +/− 6% of alleles were observed recurrently, whereas in pairwise comparisons of embryos only 2% +/− 5% of alleles were observed recurrently. The abundances of alleles were well-correlated between technical replicates for each of two 72 hpf embryos (fig. S8A and B), and alleles containing many edits were more likely to be unique to an embryo than those with few edits (fig. S8C). To assess when editing begins, we analyzed the proportions of the most common editing events across all barcodes sequenced in a given embryo, reasoning that the earliest edits would be the most frequent. Across eight v6 and 45 v7 embryos, we never observed an edit that was present in 100% of cells. This observation indicates that no permanent edits were introduced at the one-cell stage. In nearly all embryos, we observe that the most common edit is present in >10% of cells, and in some cases in ~50% of cells (Fig. 3D and fig. S9). This observation also holds in ~4,000-cell dome stage embryos, which result from approximately 12 rounds of largely synchronous division unaccompanied by cell death. Most of these edits are rare or absent in other embryos, suggesting they are unlikely to have arisen recurrently within each lineage. These results suggest that the edits present in ~50% of cells were introduced at the two-cell stage and that the edits present in >10% of cells were introduced before the 16-cell stage.

How long does barcode editing persist? Two aspects of the data suggest that it tapers relatively early in development. First, in dome stage embryos (4.3 hpf), we captured barcodes from a median of 2,086 cells, in which a median of 4.8 targets were edited. Although the number of cells and alleles that we were able to sample increased at the later developmental stages, the proportion of edited sites appeared relatively stable (Fig. 3C). If editing were occurring throughout this time course, we would instead expect the proportion of edited sites to increase substantially. Second, the number of unique alleles appears to saturate early, never exceeding 4,200 (Fig. 3E). For example, only 4,195 alleles were observed in a 72 hpf embryo in which we sampled the highest number of cells (n = 31,639). These results suggest that the majority of editing events occurred before dome stage.

### Editing diversity in adult organs

To evaluate whether barcodes edited during embryogenesis can be recovered in adults, we dissected two edited 4-month old v7 transgenic zebrafish (ADR1 and ADR2) (Fig. 5A). We collected organs representing all germ layers - the brain and both eyes (ectodermal), the intestinal bulb and posterior intestine (endodermal), the heart and blood (mesodermal), and the gills (neural crest, with contributions from other germ layers). We further divided the heart into four samples – a piece of heart tissue, dissociated unsorted cells (DHCs), FACS-sorted GFP+ cardiomyocytes, and non-cardiomyocyte heart cells (NCs) (fig. S10). We isolated genomic DNA from each sample, amplified and sequenced edited barcodes with high technical reproducibility (fig. S11), and observed barcode editing rates akin to those in embryos (fig. S12). For zebrafish ADR1, we captured barcodes from between 776 and 44,239 cells from each tissue sample (median 17,335), corresponding to a total of 197,461 cells and 1,138 alleles. For zebrafish ADR2, we captured barcodes from between 84 and 52,984 cells from each tissue sample (median 20,973), corresponding to a total of 217,763 cells and 2,016 alleles. These results show that edits introduced to the barcode during embryogenesis are inherited through development and tissue homeostasis and can be detected in adult organs.

**Fig. 5.**
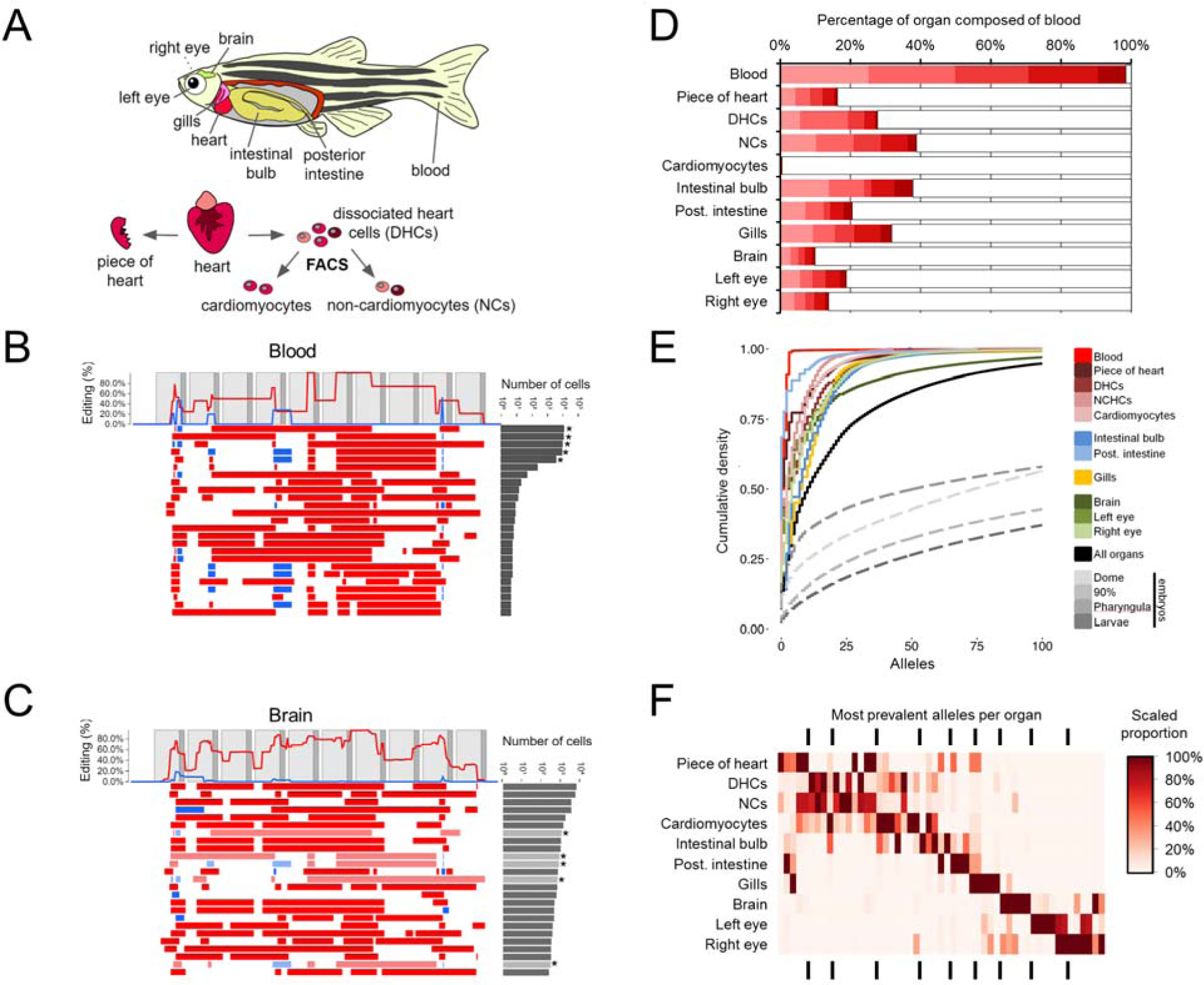
Organ-specific progenitor cell dominance. (**A**) The indicated organs were dissecte**d** from a single adult v7 transgenic edited zebrafish (ADR1). A blood sample was collected as described in the Methods. The heart was further split into the four samples shown (fig. S10). (**B**) Patterns of editing in the most prevalent 25 alleles (out of 135 total) recovered from the blood sample. Layout as described in the Fig. 1B legend. The most prevalent 5 alleles (indicated by asterisks) comprise >98% of observed cells. (**C**) Patterns of editing in the most prevalent 25 alleles (out of 399 total) recovered from brain. Layout as described in the Fig. 1B legend. Alleles that are identical in sequence to the most prevalent blood alleles are indicated by asterisks and light shading. (**D**) The five dominant blood alleles (shades of red) are present in varying proportions (10-40%) in all intact organs except the FACS-sorted cardiomyocyte population (0.5%). All other alleles are summed in grey. (**E**) The cumulative proportion of cells (y-axis) represented by the most frequent alleles (x-axis) for each adult organ of ADR1 is shown, as well as the adult organs in aggregate. In all adult organs except blood, the five dominant blood alleles are excluded. All organs exhibit dominance of sampled cells by a small number of progenitors, with fewer than 7 alleles comprising the majority of cells. For comparison, a similar plot for the median embryo (dashed) from each time-point of the developmental time course experiment is also shown. (**F**) The distribution of the most prevalent alleles for each organ, after removal of the five dominant blood alleles, across all organs. The most prevalent alleles were defined as being at >5% abundance in a given organ (median 5 alleles, range 4-7). Organ proportions were normalized by column and colored as shown in legend. Underlying data presented in table S2.

### Differential contribution of embryonic progenitors to adult organs

To analyze the contribution of diverse alleles to different organs, we compared the frequency of edited barcodes within and between organs. We first examined blood (of note, zebrafish erythrocytes are nucleated (30)). Only 5 alleles defined over 98% of cells in the ADR1 blood sample (Fig. 5B), suggesting highly clonal origins of the adult zebrafish blood system from a few embryonic progenitors. Consistent with the presence of blood in all dissected organs, these common blood alleles were also observed in all organs (10-40%; Fig. 5C) but largely absent from cardiomyocytes isolated by flow sorting (0.5%). Furthermore, the relative proportions of these five alleles remained constant in all dissected organs, suggesting that they primarily mark the blood and do not substantially contribute to non-blood lineages (Fig. 5D). In performing similar analyses of clonality across all organs (while excluding the five most common blood alleles), we observed that a small subset of alleles dominates each organ (Fig. 5E). Indeed, for all dissected organs, fewer than 7 alleles comprised >50% of cells (median 4, range 2-6), and, with the exception of the brain, fewer than 25 alleles comprised >90% of cells (median 19, range 4-38). Most of these dominant alleles were organ-specific, *i.e*. although they were found rarely in other organs, they tended to be dominant in only one organ (Fig. 5F). For example, the most frequent allele observed in the intestinal bulb comprised 13.6% of captured non-blood cells observed in that organ, but <0.01% of cells observed in any other organ. There are exceptions, however. For example, one allele is observed in 24.7% of sorted cardiomyocytes, 13.4% of the intestinal bulb, and at lower abundances in all other organs. Similar results were observed in ADR2 (fig. S13). These results indicate that the majority of cells in diverse adult organs are descended from a few differentially edited embryonic precursors.

### Reconstructing lineage relationships in adult organs

To reconstruct the lineage relationships between cells both within and across organs on the basis of shared edits, we again relied on maximum parsimony methods (fig. S4B). The resulting trees for ADR1 and ADR2 are shown in Fig. 6 and fig. S14, respectively. We observed clades of alleles that shared specific edits. For example, ADR1 had 8 major clades, each defined by ‘ancestral’ edits that are shared by all captured cells assigned to that clade (Fig. 7A; also indicated by colors in the tree shown in Fig. 6). Collectively, these clades comprised 49% of alleles and 90% of the 197,461 cells sampled from ADR1 (Fig. 7A). Blood was contributed to by 3 major clades (#3, #6, #7) (Fig. 7B). After re-allocating the 5 dominant blood alleles from the composition of individual organs back to blood (Fig. 5B and fig. S15), we observed that all major clades made highly non-uniform contributions across organs. For example, clade #3 contributed almost exclusively to mesodermal and endodermal organs, while clade #5 contributed almost exclusively to ectodermal organs. These results reveal that GESTALT can be used to infer the contributions of inferred ancestral progenitors to adult organs.

**Fig. 6.**
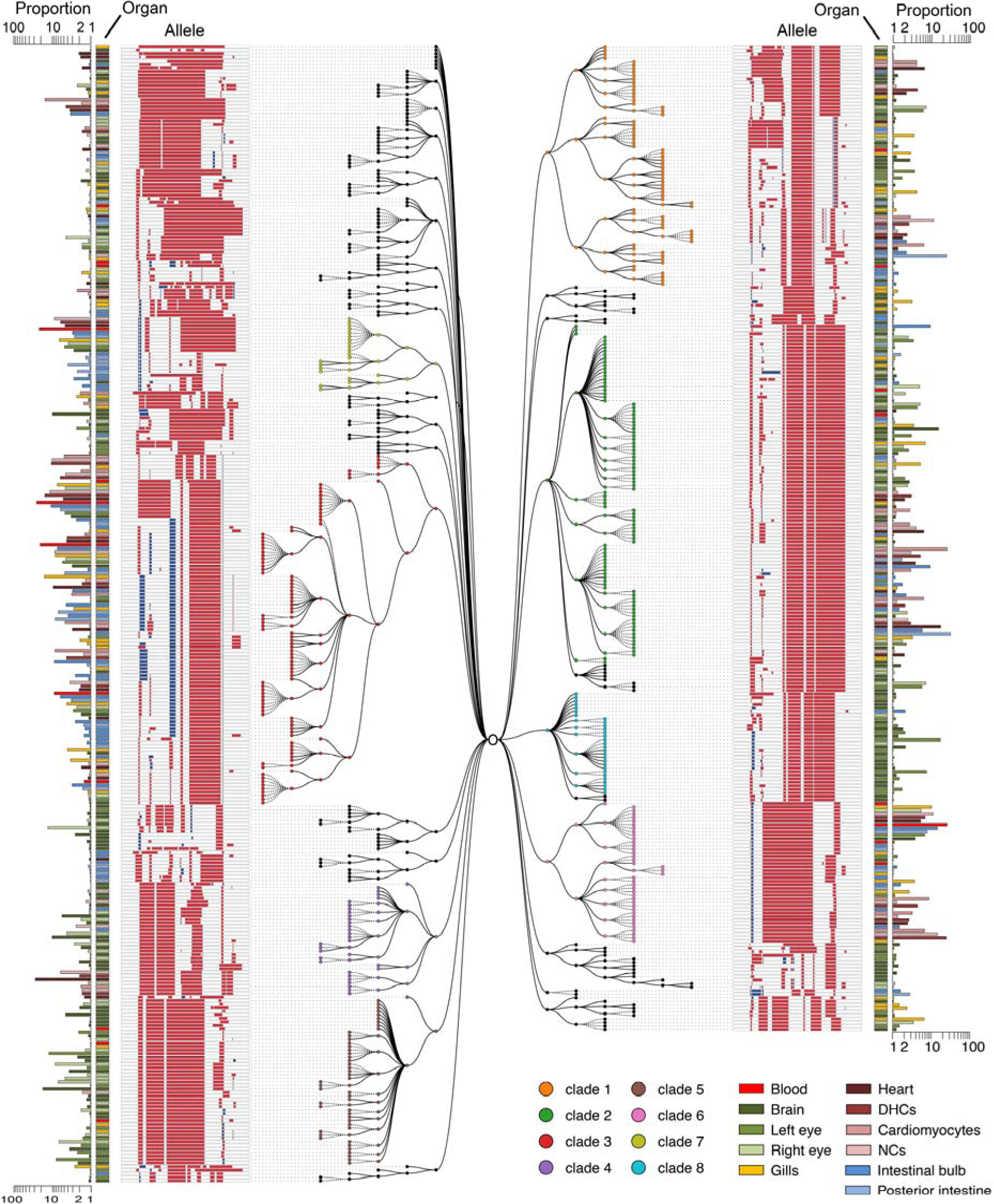
Lineage reconstruction for adult zebrafish ADR1. Unique alleles sequenced from adult zebrafish organs can be related to one another using a maximum parsimony approach into a multifurcating lineage tree. For reasons of space, we show a tree reconstructed from the 611 ADR1 alleles observed at least 5 times in individual organs. Eight major clades are displayed with colored nodes, each defined by ‘ancestral’ edits that are shared by all alleles assigned to that clade (shown in Fig. 7A). Editing patterns in individual alleles are represented as shown previously. Alleles in multiple organs are plotted on separate lines per organ and these nodes connected with stippled branches. Two sets of bars outside the alleles identify the organ in which the allele was observed and the proportion of cells in that organ represented by that allele (log scale).

**Fig. 7.**
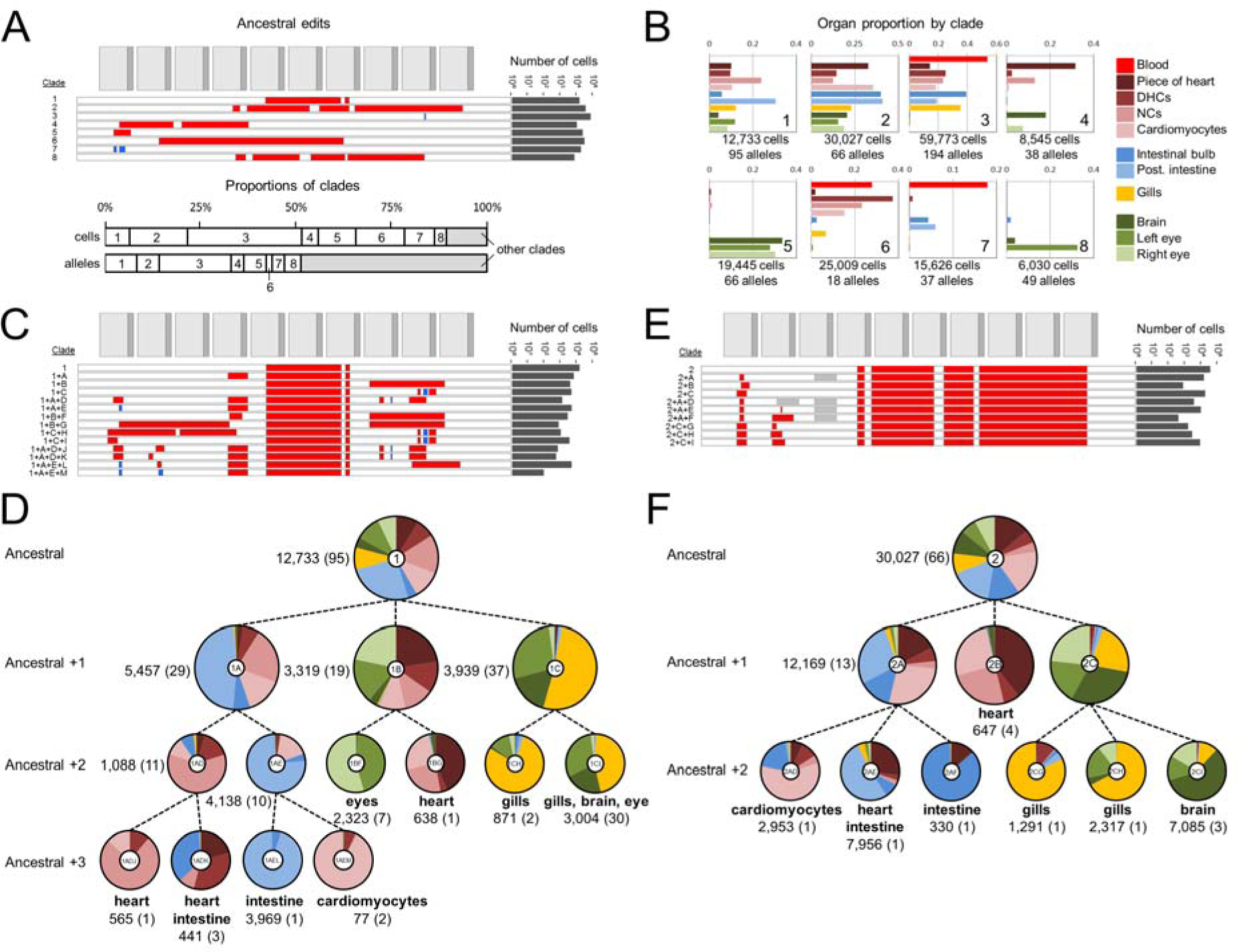
Clades and subclades corresponding to inferred progenitors exhibit increasing levels of organ restriction. (**A**) Top panel: The inferred ancestral edits that define eight major clades of ADR1, as determined by parsimony, are shown, with the total number of cells in which these are observed indicated on the right. Bottom panel: Contributions of the eight major clades to all cells or all alleles. 19 alleles (out of 1,138 total) that contained ancestral edits from more than one clade were excluded from assignment to any clade, and any further lineage analysis. (**B**) Contributions of each of the eight major clades to each organ, displayed as a proportion of each organ. To accurately display the contributions of the eight major clades to each organ, we first re-assigned the five dominant blood alleles from other organs back to the blood. The total number of cells and alleles within a given major clade are listed below. The clade contributions of all clades and subclades are presented in table S3. For heart subsamples, ‘piece of heart’ = a piece of heart tissue, ‘DHCs’ = dissociated unsorted cells; ‘cardiomyocytes’ = FACS-sorted GFP+ cardiomyocytes; and ‘NCs’ = non-cardiomyocyte heart cells. (**C**) and (**E**) Edits that define subclades of clade #1 (C) and clade #2 (E), with the total number of cells in which these are observed indicated on the right. A grey box indicates an unedited site or sites, distinguishing it from related alleles that contain an edit at this location. (**D**) and (**F**) Lineage trees corresponding to subclades of clade #1 (D) and clade #2 (F) that show how dependent edits are associated with increasing lineage restriction. The pie chart at each node indicates the organ distribution within a clade or subclade. Ratios of cell proportions are plotted, a normalization which accounts for differential depth of sampling between organs. Labels in the center of each pie chart correspond to the subclade labels in (c/e). Alleles present in a clade but not assigned to a descendent subclade (either they have no additional lineage restriction or are at low abundance) are not plotted for clarity. The number of cells (and the number of unique alleles) are also listed, and terminal nodes also list major organ restriction(s), *i.e.* those comprising >25% of a subclade by proportion.

Although some ancestral clades appear to contribute to all germ layers, we find that subclades, defined by additional shared edits within a clade, exhibit greater specificity. For example, while clade #1 contributes substantially to all organs except blood, additional edits divide clade #1 into three subclades with greater tissue restriction (Fig. 7C and D). The #1+A subclade primarily contributes to mesendodermal organs (heart, both gastrointestinal organs) while the #1+C subclade primarily contributes to neuroectodermal organs (brain, left eye, and gills). Similar patterns are observed for clade #2 (Fig. 7E and F), where the #2+A subclade contributes primarily to mesendodermal organs, the #2+B subclade to the heart, and the #2+C clade to neuroectodermal organs. Additional edits divide these subclades into further tissue-specific sub-subclades. For example, while the #2+A subclade is predominantly mesendoderm, additional edits define #2+A+D (heart, primarily cardiomyocytes), #2+A+E (heart and posterior intestine), and #2+A+F (intestinal bulb). All of the major clades exhibit similar patterns of increasing restriction with additional edits (Fig. 7C-F and fig. S16). Similar observations were made in fish ADR2 (fig. S17). These results indicate that GESTALT can record lineage relationships across many cell divisions and capture information both before and during tissue restriction.

## Discussion

We describe a new method, GESTALT, which uses combinatorial and cumulative genome editing to record cell lineage information in a highly multiplexed fashion. We successfully applied this method to both artificial lineages (cell culture) as well as to whole organisms (zebrafish).

The strengths of GESTALT include: 1) the combinatorial diversity of mutations that can be generated within a dense array of CRISPR/Cas9 target sites; 2) the potential for informative mutations to accumulate across many cell divisions and throughout an organism’s developmental history; 3) the ability to scalably query lineage information from at least hundreds of thousands of cells and with a single sequencing read per single cell; 4) the likely applicability of GESTALT to any organism, from bacteria and plants to vertebrates, that allows genome editing, as well as human cells (*e.g.* tumor xenografts). Even in organisms in which transgenesis is not established, lineage tracing by genome editing may be feasible by expressing editing reagents to densely mutate an endogenous, non-essential genomic sequence.

Our experiments also highlight several remaining technical challenges. Chief amongst these are: 1) the chance recurrence of identical edits or similar patterns of edits in distantly related cells can confound lineage inference; 2) non-uniform editing efficiencies and inter-target deletions within the barcode contribute to suboptimal sequence diversity and loss of information, respectively; 3) the transient means by which Cas9 and sgRNAs are introduced likely restrict editing to early embryogenesis; 4) the computational challenge of precisely defining the multiple editing events that give rise to different alleles complicates the unequivocal reconstruction of lineage trees; and 5) the difficulty of isolating tissues without contamination by blood and other cells can hinder the assignment of alleles to specific organs. A broader set of challenges includes the lack of information about the precise anatomical location and exact cell type of each queried cell, the fact that genome editing events are not directly coupled to the cell cycle, and the failure to recover all cells. These challenges currently hinder the reconstruction of a lineage tree as complete and precise as the one that Sulston and colleagues described for *C. elegans*. Despite these limitations, our proof-of-principle study shows that GESTALT can inform developmental biology by richly defining lineage relationships among vast numbers of cells recovered from an organism.

The current challenges highlight the need for further optimization of the design of targets and arrays, as well as the delivery of editing reagents. For example, an array containing twice as many targets as used here could fit within a single read on contemporary sequencing platforms, thus yielding more lineage information per cell without sacrificing throughput. Also, as we have shown, adjustments to the target sequences and dosages of editing reagents can be used to fine-tune mutation rates and to minimize undesirable inter-target deletions. Finally, sgRNA sequences and lengths (31), Cas9 cleavage activity and target preferences (32, 33), and the means by which Cas9 and sgRNA(s) are expressed (e.g. transient, constitutive (34), or induced (35, 36)), can be altered to control the pace, temporal window and tissue(s) at which the barcodes are mutated. For example, coupling editing to cell cycle progression might enable higher resolution reconstruction of lineage relationships throughout development.

Our application of GESTALT to a vertebrate model organism, zebrafish, demonstrates its potential to yield insights into developmental biology. First, our results suggest that relatively few embryonic progenitor cells give rise to the majority of cells of many adult zebrafish organs, reminiscent of clonal dominance (37, 38). For example, only 5 of the 1,138 alleles observed in ADR1 gave rise to >98% of blood cells, and for all dissected organs, fewer than 7 alleles comprised >50% of cells. There are several mechanisms by which such dominance can emerge, *e.g.* by uneven starting populations in the embryo, drift, competition, interference, unequal cell proliferation or death, or a combination of these mechanisms (39-42). Controlling the temporal and spatial induction of edits and isolating defined cell types from diverse organs should help resolve the mechanisms by which different embryonic progenitors come to dominate different adult organs.

Second, we show that GESTALT can inform the lineage relationships amongst thousands of differentiated cells. For example, following the accumulation of edits from ancestral to more complex reveals the progressive restriction of progenitors to germ layers and then organs. Cells within an organ can both share and differ in their alleles, revealing additional information about organ development. Future studies will need to determine whether such lineages reflect distinct cell fates (*e.g.*, blood sub-lineages or neuronal subpopulations), because the anatomical resolution at which we queried alleles was restricted to grossly dissected organs and tissues. Because edited barcodes are expressed as RNA, we envision that combining our system with other platforms will permit much greater levels of anatomical resolution without sacrificing throughput. For example, *in situ* RNA sequencing of barcodes would provide explicit spatial and histological context to lineage reconstructions (19, 20). Also, capturing richly informative lineage markers in single cell RNA-seq or ATAC-seq datasets may inform the interpretation of those molecular phenotypes, while also adding cell type resolution to studies of lineage (43, 44). Such integration may be particularly relevant to efforts to build comprehensive atlases of cell types. Because these single cell methods generate many reads per single cell, this would also facilitate using multiple, unlinked target arrays. In principle, the combined diversity of the barcodes queried from single cells could be engineered to uniquely identify every cell in a complex organism. In addition, orthogonal imaging-based lineage tracing approaches in fixed and live samples (*e.g.*, Brainbow and related methods (16, 29)) and longitudinal whole animal imaging approaches (45, 46) might be leveraged in parallel to validate and complement lineages resolved by GESTALT.

Although further work is required to optimize GESTALT towards enabling spatiotemporally complete maps of cell lineage, our proof-of-principle experiments show that using multiplex *in vivo* genome editing to record lineage information to a compact barcode at an organism-wide scale will be a powerful tool for developmental biology. This approach is not limited to normal development but can also be applied to animal models of developmental disorders, as well as to investigate the origins and progression of cancer. Our study also supports the notion that whereas its most widespread application has been to modify endogenous biological circuits, genome editing can also be used to stably record biological information (47), analogous to recombinase-based memories but with considerably greater flexibility and scalability. For example, coupling editing activity to external stimuli or physiological changes could record the history of exposure to intrinsic or extrinsic signals. In the long term, we envision that rich, systematically generated maps of organismal development, wherein lineage, epigenetic, transcriptional and positional information are concurrently captured at single cell resolution, will advance our understanding of normal development, inherited diseases, and cancer.

## Author contributions

GF, AM and JS developed the initial concept. GF led the cell culture experiments and developed the UMI protocol, with assistance from AM. JG led the fish experiments, with assistance from AM and GF. AM led development of the analysis pipeline. AM, GF, and JG processed and analyzed the data. AM, GF, JG, AS and JS designed experiments and interpreted the data. MH provided critical early insight. AM, GF, JG, AS, and JS wrote the manuscript.

## Materials and Methods

### Design of synthetic target arrays

Barcodes were designed as arrays of nine to twelve sense-oriented CRISPR/Cas9 target sites (23 bp protospacer plus PAM sequences) separated by 3-5 bp linker sequences. Four initial designs (barcodes v1-v4) comprised of target sites for the sgRNA spacer sequence: 5’-GGCACTGCGGCTGGAGGTGG. The v1 barcode was comprised of ten targets arrayed in order of decreasing activity as measured with the GUIDE-seq assay performed in human cells (*22*), starting with the target perfectly matching the sgRNA spacer sequence. The v2-v4 barcodes comprised of nine to ten non-overlapping target sets, all with activities less than half the perfectly matching target in the GUIDE-seq assay. To reduce repetitive subsequences within each barcode, protospacers were chosen such that no 8 bp sequence was present in more than one protospacer within each barcode. After testing activities of targets in the v1-v4 barcodes in cell culture, the v5 barcode was designed to contain twelve targets that showed greater than ~1% editing activity, including v1 targets 1-6, v3 target 1, v2 targets 1, 2 and 5, and v4 targets 1 and 3.

Two new barcodes, v6 and v7, were designed for use in zebrafish, each with ten CRISPR target sites not found in the *D. rerio* genome. Candidate target sequences were screened to remove any homopolymer runs, outside of the NGG of the protospacer, and were selected for editing activity [http://crispr.mit.edu]. The v6 and v7 barcodes were constructed as a series of 10 protospacer sequences meeting these criteria, with 4 bp linkers.

Each barcode was ordered as a gBlock (IDT) with ends compatible for In-Fusion cloning (ClonTech) into the 3’ UTR of the EGFP gene in the lentiviral construct pLJM1-EGFP (Addgene #19319).

The sequences of all barcodes (v1 through v7) are provided in table S4.

### Generation of cell lines containing synthetic target arrays

To generate cell lines harboring single copies of barcodes, lentiviral particles were produced in HEK 293T cells transfected with lentivirus V2 packaging plasmids and barcode constructs. Viral supernatant harvested three days post transfection was used at low MOI to transduce 293T cells (MOI < 0.2). Successfully transduced cells were selected using puromycin (2 ug/ml), yielding polyclonal, barcode+ populations for barcodes v1-v5. Three monoclonal lines each harboring barcode v1 were generated by single-cell FACS, and used experimentally to compare editing rates across different integration sites. One of these was used as the parent line for cell culture lineages derived using barcode v1.

### Editing of barcodes in cell lines

293T populations bearing barcodes v1-v5 were grown to 50-90% confluency in a 6-well dish. Cells were co-transfected using Lipofectamine 3000 (Life Technologies) according to protocol with 2 μg pX330-v1 and 0.5 μg pDsRed in a 6-well dish. One to three days post transfection, the cells were sorted on an Aria III FACS machine for DsRed fluorescence (as a marker transfection). As indicated, either DsRed low, DsRed high, or total DsRed populations were sorted and cultured. At 7 days post-transfection, cells were harvested for gDNA preparation using the Qiagen DNeasy kit.

To stably deliver Cas9 and the sgRNA via lentivirus, the spacer sequence was cloned into the plasmid LentiCrispr v2 (Zhang lab, Addgene #52961) and virus was produced in 293T cells in the same manner described above. Wild-type 293T cells were transduced with pLenti-Crispr-V2-HMID.v1 and selected with puromycin, and then transduced with lentivirus bearing barcode v5. To impose a bottleneck, 200 GFP+ cells were sorted from this population and expanded under puromycin selection for two weeks prior to sampling gDNA.

### Cell culture lineage experiments

Twelve lineages were established from a monoclonal barcode v1 293T cell line by transfecting cells as described above, and sorting single DsRed-low cells into a 96-well plate (DsRed low cells were used to limit Cas9 delivery and thus potential saturation of possible edits in this initial editing round). Cell sorting was performed seven days post-transfection, to reduce the likelihood that additional edits would arise after lineages were separated. Single cell-derived populations were expanded in culture for 3 weeks. A sample of cells from each lineage was pelleted and frozen. Next, each of the twelve lineages were transfected a second time, to induce another round of editing. Two 100-cell DsRed-low populations from each lineage were sorted 4-days post-transfection, and cultured to confluence in 96-well plates before harvesting gDNA.

Four additional monoclonal populations bearing v5 barcodes edited via transfection of pX330-v1 were also isolated by single-cell sorting. Re-editing of each population was achieved by two successive rounds of transfection with pX330-v1 (3 days apart). Cells were harvested for gDNA one week after the second transfection.

### Barcode amplification and sequencing protocols

Kapa High Fidelity Polymerase was used for all barcode amplification steps. Gradient PCRs were performed to optimize annealing temperatures for amplification from gDNA. For experiments performed without UMIs, up to 250 ng of gDNA was loaded into a single 50 μl PCR reaction and amplified using primers immediately flanking the barcode (see table S4 for oligo sequences). If there was less than 250 ng from a sample, all of it was used in a single reaction. For experiments performed with UMIs, a primer with a sequencing adapter and 10 nt of fully degenerate sequence 5’ to the barcode-flanking sequence was used for a single prolonged extension step, in which the temperature was ramped between annealing and extending for five cycles (without a denaturing step to prevent re-sampling of gDNA barcodes). All cell culture experiments and v6 zebrafish embryos received a single extension to incorporate UMIs, whereas v7 embryo time-course experiments and all ADR1 tissues (also v7) received 2 UMI incorporation cycles due to having low gDNA consequent to fewer cells being present in early embryo and sorted heart samples. To minimize repetitive amplification of the same barcode, no reverse primer was included in UMI-tagging reactions. DNA was then purified using AMPure beads (Agencourt), and loaded into a PCR primed from the sequencing adapter flanking each UMI and a site immediately 3’ of the barcode.

For all experiments, two ensuing qPCRs were performed prior to sequencing to incorporate sequencing adapters, sample indexes, and flow cell adapters. AMPure beads were used to purify PCR products after each reaction.

Paired-end sequencing was performed on an Illumina MiSeq using 500- or 600-cycle kits for all cell culture experiments. Zebrafish experiments were sequenced on an Illumina NextSeq using 300-cycle kits. All sequencing generated adequate depth to sample each barcode present in a given sample to an average of greater than 10x coverage. To minimize contributions from sequencing error a read threshold was used for calling unique barcodes. This was conservatively set by dividing the number of reads from a sample by the number of expected barcode copies to be present in the amount of gDNA loaded into each PCR based on the assumption that each cell contributed a single barcode.

Sequencing data for all samples was processed in a custom pipeline available on GitHub (https://github.com/shendurelab/Cas9FateMapping). Briefly, amplicon sequencing reads were first processed with the Trimmomatic software package to remove low quality bases (fig. S4A)(48). The resulting reads were then grouped by their UMI tag. A raw read count threshold was set for each experiment based on sequencing depth, such that only UMIs observed in at least that many reads were analyzed to minimize contributions from sequencing error. For each UMI, a consensus sequence was called by jointly aligning all UMI-matched reads using the MAFFT multiple sequence aligner. These reads were merged using the FLASH (49) read merging tool, and both merged and unmerged reads were aligned to the amplicon reference using the NEEDLEALL (50) aligner with a gap open penalty of 10 and a gap extension penalty of 0.5. To capture read-through, UMI degenerate bases and adapter sequences were included in the reference amplicon sequence, and mismatches to Ns in the degenerate bases were set to a penalty of 0. To eliminate off-target sequencing reads, aligned sequences were required to match greater than 85% of bases at non-indel positions, to have correct PCR primer sequences on both the 5’ and 3’ ends, and to match at least 50 bases of the reference sequence (including primer sequences). Target sites were deemed edited if there was an insertion or deletion event present within 3 bases of the predicted Cas9 cut site (3 nucleotides 5’ of each PAM), or if a deletion spanned the site entirely. Sites were marked as disrupted if there was not perfect alignment of the barcode over the entirety of the reference target sequence. An edited barcode was then defined as the complete list of insertion and deletion events (*i.e.* ‘editing events’) within the consensus sequence for a given UMI.

### Maximum parsimony lineage reconstruction

For lineage reconstruction (fig. S4B), recurrently observed barcode alleles within a single organ or cell population were reduced to a single representative entry. We then used Camin-Sokal maximum parsimony to reconstruct lineages, as implemented in the PHYLIP MIX software package (26). Camin-Sokal maximum parsimony assumes that the initial cell or zygote is unedited, and that editing is irreversible. To run MIX, a matrix was created where each row corresponded to an allele, and each column corresponds to a unique editing event. Each entry in this matrix is an indicator variable of presence or absence of a specific edit in that allele (1 or 0). Events were also weighted by their log-abundance and scaled to the range allowed in MIX (0-Z). MIX was run, and the output was parsed to recover edit patterns in ancestral nodes. Internal parent-child nodes that had identical editing patterns were collapsed using the recovered ancestral alleles and the output tree. When a parent node and child node share the same allele, the grandchildren nodes were transferred to the parent and the child node was removed, creating multifurcating parent nodes. The resulting tree was converted to an annotated JSON tree compatible with our visualization tools. All code is available on the Shendure lab github website: https://github.com/shendurelab/Cas9FateMapping.

### Zebrafish husbandry

All vertebrate animal work was performed at the facilities of Harvard University, Faculty of Arts & Sciences (HU/FAS). This study was approved by the Harvard University/Faculty of Arts & Sciences Standing Committee on the Use of Animals in Research & Teaching under Protocol No. 25–08. The HU/FAS animal care and use program maintains full AAALAC accreditation, is assured with OLAW (A3593-01), and is currently registered with the USDA.

### Cloning transgenesis vector

The transgenesis vectors pTol2-DRv6 and pTol2-DRv7 were constructed as follows. The v6 or v7 array was cloned into the 3’ UTR of a DsRed coding sequence under control of the ubiquitin promoter (51). This cassette was placed in a Tol2 transgenesis vector containing a cmlc2:GFP marker, which drives expression of GFP in the cardiomyocytes of the heart from 24 hpf to adulthood (52). Plasmids are available from Addgene.

### Generating transgenic zebrafish

To generate founder fish, 1-cell embryos were injected with zebrafish codon optimized Tol2 mRNA and pTol2-DR1v6 or pTol2-DR1v7 vector. Potential founder fish were screened for heart GFP expression at 30 hpf and grown to adulthood. Adult founder transgenic fish were identified by outcrossing to wild type and screening clutches of embryos for heart GFP expression at 30 hpf.

### Transgene copy number quantification

To identify single copy Tol2 transgenics, copy number was quantified using qPCR (*29*). Briefly, genomic DNA was extracted from candidate embryos or fin-clips of adult fish using the HotSHOT method (53) and subjected to qPCR using a set of primers targeting DsRed and a set targeting a diploid conserved region of the genome (table S4) and compared to reference non-transgenic, 1-copy and 2-copy transgenic animals using the ddCt method.

### Generation and delivery of editing reagents

sgRNAs specific to each site of the v6 or v7 array were generated as previously described (54), except that sgRNAs were isolated after transcription by column purification (Zymogen). 1-cell embryos resulting from an outcross of a transgenic founder were injected with two different volumes (0.5 nl, 1/3x or 1.5nl, 1x) of Cas9 protein (NEB) and sgRNAs in salt solution (8 μM Cas9, 100 ng/μl pooled sgRNAs, 50 mM KCl, 3 mM MgCl_2_, 5 mM Tris HCl pH 8.0, 0.05% phenol red). Transgenic embryos were collected at the time points indicated in the text and genomic DNA extracted as described below. To confirm editing, PCR was conducted on a subset of samples using primers flanking the v6 or v7 array (table S4), and amplicons were loaded on a 2% agarose gel for electrophoresis.

### Imaging

Embryos were anaesthetized and manually dechorionated in MS222, mounted in methylcellulose and imaged using a Leica upright fluorescence microscope.

### Organ Dissection

Adult edited single copy transgenic fish were isolated without food for one day to reduce food particles in the gastrointestinal system, then anaesthetized in MS222 and euthanized on ice. Before dissection, blood was collected using a centrifugation method (55). This collection method greatly enriches for blood cells, particularly red blood cells, but also results in contamination from skin or other tissues. The fish were pinned on a silicon mat and surgery was conducted using sterile tools to remove organs as in (56). Organs were washed in PBS and, with the exception of the heart, frozen in tubes on dry ice. A piece of heart tissue was collected before the remainder of the heart was dissociated following manufacturer's instructions (Miltenyi # 130-098-373). After dissociation, a sample of dissociated heart cells was collected (DHCs), and the remaining cells sorted using a Beckman Coulter MoFlo XDP Cell Sorter through a series of three gates to minimize debris and cell doublets, and then split into two additional populations: GFP+ cardiomyocytes and GFP-non-cardiomyocyte heart cells (NCs, fig. S10).

### Genomic DNA preparation from zebrafish embryos and organs

Zebrafish embryo and adult organ gDNA was prepared using the Qiagen DNeasy kit. For heart samples from cell sorting experiments, 1 μl of poly-dT carrier DNA (25 uM) was added prior to gDNA preparation. Digestion with proteinase K at 56° C was performed overnight for intact organs (brain, eyes, gills, intestinal bulb, posterior intestine, and piece of heart) and for 30 minutes for blood samples, dissociated heart cells and embryos. gDNA was eluted in 100 μl, then concentrated using an Eppendorf Vacufuge for samples yielding less than 1 μg.

## Acknowledgements

For discussion and advice, we thank Joe Felsenstein, Mary Kuhner, Marlies Rossmann, Eva Fast, Caroline Burns, Raz Ben-Yair, Steve Salipante; members of the Shendure Lab, particularly Matthew Snyder; members of the Schier Lab, particularly Jeffrey Farrell, Bushra Raj, Megan Norris, and Nathan Lord; and members of the Horwitz lab, particularly Donovan Anderson. We thank George Church for work on predecessors of this concept with JS in 2000. We thank Michael Desai, Rich Losick, Andrew Murray and Len Zon for comments on the manuscript. We also thank the Bauer Core FACS Facility, the staff of the zebrafish facility, and Lindsey Pieper for technical support. This work was supported by grants from the Paul G. Allen Family Foundation (JS & MSH), an NIH Director’s Pioneer Award (JS; DP1HG007811), NIH/NIGMS (AFS; GM056211), NIH/NICHD (AFS; HD085905), and NIH/NIMH (AFS; MH105960). JAG was supported by a fellowship from the American Cancer Society. AHM was supported by a fellowship from the NIH/NHLBI (T32HL007312). JS is an investigator of the Howard Hughes Medical Institute.

